# Blocking primer improves detection of tick-borne pathogens in *Ixodes scapularis* (black-legged ticks) from a Lyme disease hotspot region in eastern Ontario, Canada

**DOI:** 10.64898/2026.08.17.745364

**Authors:** Sreevatshan K. Srinivasan, Sima Afsharnezhad, Amber R. Paulson, Damian C. Bourne, Zhengxin Sun, Lisa F. Carver, Jad Tirani, Henry Wong, Calvin J. Sjaarda, Shu He, Prameet M. Sheth, Robert I. Colautti

**Affiliations:** Department of Biology, Queen’s University, 116 Barrie Street, Kingston, Ontario, K7L 3N6, Canada; Environmental Assessment Office, British Columbia Ministry of Environment and Parks, Victoria, BC, V8W 9V1, Canada; Department of Biomedical and Molecular Sciences, Queen’s University, 18 Stuart St., Kingston, ON, K7L 3N6, Canada; Department of Pathology and Molecular Medicine, Queen’s University, 88 Stuart St., Kingston, ON, K7L 2V6 Canada; Division of Microbiology, Kingston Health Sciences Center, 76 Stuart St., Kingston, ON, K7L 2V7 Canada

## Abstract

Tick-borne pathogen (TBP) surveillance strategies that rely exclusively on targeted methods like PCR (PCR) or immunoblots do not benefit from strain-level sequence variation. Bacterial 16S rRNA metabarcoding offers more agnostic detection but is constrained in *I. scapularis* by the dominance of a maternally inherited endosymbiont, *Rickettsia buchneri*. Here we report the design and evaluation of three *R. buchneri*-specific blocking primers to suppress endosymbiont amplification during full-length 16S rRNA library preparation. Of these, primer 18F-Rb-C3 reduced *R. buchneri* relative abundance approximately 32-fold. We applied 18F-Rb-C3 with V4-16S metabarcode sequencing on 67 ticks collected from farm animals in Eastern Ontario and compared *Borrelia* species detection against qPCR. The V4-16S rRNA metabarcoding identified *Borrelia* species in 21 samples, whereas qPCR detected *Borrelia* in 24 samples and 11 samples were detected by both methods. Additionally, metabarcoding detected *Anaplasma phagocytophilum* in 12 samples, including seven samples coinfected with *Borrelia,* in the same assay. Variation relevant to strain surveillance was also detected by sequencing, though V4-16S was not sufficient to resolve closely related *Borrelia* genospecies or *A. phagocytophilum* variants. These findings demonstrate that blocking primer 18F-Rb-C3 enhances sensitivity of amplicon sequencing to the level of qPCR while also detecting other pathogens and sequence variants in a single assay.

## 1 Introduction

Ticks in the family Ixodidae transmit over 40% of the emerging vector-borne diseases, imposing a significant and growing burden on public healthcare systems (Swei et al. 2020; Rochlin and Toledo 2020). In Canada, *Ixodes scapularis* (black-legged ticks) has been rapidly expanding its geographic range since the 1970s and acts as a vector for diseases such as Lyme disease (*Borrelia burgdorferi sensu lato* (Bbsl) complex), anaplasmosis (*Anaplasma phagocytophilum*), babesiosis (*Babesia* spp.), and Powassan virus disease, and for emerging pathogens including *Ehrlichia* spp. (Ehrlichiosis) ((Watson and Anderson, 1976; Bouchard et al. 2019; Crandall et al. 2024; Westcott et al. 2025). Anthropogenic effects, including habitat fragmentation and climate change, have resulted in shifts in species distributions, contributing to changes in tick-borne disease dynamics, including increased tick prevalence and the distribution of tick-borne pathogens (TBPs) at an unprecedented rate (Diuk-Wasser et al. 2021; Ronald Rosenberg et al. 2018). Therefore, to understand the prevalence and dynamics of TBPs, broad-range surveillance is essential, as it will improve disease dynamics modeling and the prediction of TBPs emergence.

At present, surveillance and monitoring efforts by public health agencies to detect bacterial TBPs rely on targeted qPCR (quantitative Polymerase Chain Reaction), which is rapid, cost-effective, and sensitive (Wilson et al. 2022). However, these approaches have several limitations that constrain the detection of TBPs over a broad range. As ticks and TBPs continue to expand their range northward due to climate change, the emergence of novel bacterial TBPs in previously unaffected regions is expected. Targeted qPCR screening panels are designed for currently prevalent TBPs; they will fail to detect novel TBPs. qPCR specificity also depends on the binding affinity between probes and targets, which can be compromised by sequence variation among bacterial strains at hybridization sites, reducing binding affinity and increasing false-negative rates (Dymond 2013; Benevides Lima et al. 2022). This is particularly relevant to *Borrelia* genospecies, in which targets such as *ospA* and *ospC* are known to vary among strains and genospecies (Wallich et al. 1992; Tokarz and Lipkin 2021; Mechai et al. 2025). Furthermore, higher cycle numbers increase qPCR sensitivity but also increase the risk of off-target amplification and false positives (Dymond 2013; Benevides Lima et al. 2022). These limitations highlight the need for broad-range detection of TBPs in ticks that is agnostic, scalable, and robust to strain-level sequence variation.

High-throughput Sequencing (HTS) technologies based on amplification of the 16S rRNA gene (hereafter referred to as 16S) enable agnostic pathogen surveillance across multiple taxa in a single assay (Gu et al. 2019). Unlike qPCR primers, which are specific to a single target, universal 16S primers target highly conserved regions of the ribosomal gene, making amplification less susceptible to false negatives from strain-level sequence variation (Johnson et al. 2019). Furthermore, off-target amplicons are identified and removed during bioinformatics processing by confirming their sequences against the reference database. Previous *I. scapularis* studies using bacterial 16S metabarcoding have detected and characterized pathogens such as Bbsl, *A. phagocytophilum*, and *B. miyamotoi*, confirming that HTS can broaden the detection of bacterial TBPs (Sperling et al. 2017; Thapa et al. 2019; Price et al. 2021; Paulson et al. 2023). However, the sensitivity of 16S metabarcoding for TBP detection is constrained by sequencing depth in *I. scapularis*, which is frequently infected with the intracellular endosymbiont *Rickettsia buchneri* (Kurtti et al. 2015). *R. buchneri* is horizontally transmitted, and this dominance can reduce the proportion of sequencing reads available for pathogen taxa, thereby lowering the likelihood of detecting rare or low-abundance TBPs (Gofton, Oskam, et al. 2015; Gofton, Doggett, et al. 2015; Brinkerhoff et al. 2020). Therefore, to improve sensitivity for detecting low-abundant TBPs in *I. scapularis* using 16S metabarcoding, strategies should be developed to reduce the representation of *R. buchneri* during library preparation and/or sequencing.

One promising strategy for reducing highly abundant non-target reads is the use of blocking primers, which are primers designed against a target sequence to prevent its amplification, thereby enabling detection of low-abundance taxa without reducing sequencing depth (Gofton, Oskam, et al. 2015; Tan and Liu 2018; Gofton, Doggett, et al. 2015; Santos-Garcia et al. 2020). Currently, two mechanisms of blocking primers exist. Annealing-inhibition primers compete with the universal primers for the same binding sites, whereas elongation-arrest primers bind between the universal primers and prevent elongation (Vestheim and Jarman 2008). Studies have established that annealing-inhibiting primers block target amplification more effectively than elongation-arrest primers (Peano et al. 2005; Vestheim and Jarman 2008; Vestheim et al. 2011; Gofton, Oskam, et al. 2015). Alternative target depletion strategies, such as restriction enzyme-based targeted cleavage or PCR clamping via peptide nucleic acids (PNAs) and locked nucleic acids (LNAs), are present but limited by high costs, complexity, and lengthy synthesis times (Briones and Moreno 2012), making annealing-inhibiting blocking primers a more scalable approach for HTS.

Blocking primers designed for 16S regions have proven effective for suppressing dominant bacterial endosymbionts and improving the detection of low-abundance taxa in Arthropod microbiome studies. In ticks, an annealing-inhibiting blocking primer overlapping the 27F-Y (V1) binding region was designed to suppress “*Candidatus* Midichloria mitochondrii”, which is the maternally inherited endosymbiont of *I. holocyclus* (Gofton, Oskam, et al. 2015).This enabled detection of potential TBPs, including *Bartonella henselae*, “*Candidatus* Neoehrlichia spp.,” *Clostridium histolyticum*, and *Leptospira inadai* (Gofton, Oskam, et al. 2015). Similarly, in the whitefly (*Bemisia tabaci*), a dual-priming oligonucleotide overlapping the 515R (V3) binding region of the 16S was designed to simultaneously suppress amplicons from *Rickettsia* spp.*, Hamiltonella* spp., and *Portier* spp., which revealed previously undetected members of the microbiome (Santos-Garcia et al. 2020). However, a blocking primer targeting *Rickettsia-*derived 16S amplicon sequences from *I. scapularis* has not yet been designed to suppress *R. buchneri-*derived amplicon sequences, which motivates this study.

To improve sensitivity in detecting bacterial TBPs using HTS and enable agnostic detection beyond the constraints of targeted TBP detection, we designed three 16S blocking primer candidates and evaluated them to identify the optimal blocking primer that effectively suppresses the rickettsial endosymbiont while minimizing bias toward other bacterial taxa. We then assessed the sensitivity of the optimized 16S metabarcoding protocol by comparing it with the gold standard, targeted qPCR, using ticks collected from farm animals in Eastern Ontario. We demonstrated that the 16S metabarcoding approach, integrated with the rickettsial blocking primer, improved sensitivity in detecting bacterial TBPs in *I. scapularis*, with implications for public health surveillance.

## 2 Materials and Methods

### 2.1 Design of blocking primer

To suppress the preferential amplification of abundant *R. buchneri* reads, we developed three taxon-specific blocking primer candidates (18F-Rb-C3, 22F-Rb-C3, and 1492R-Rb-C3) for use alongside the universal full-length 16S rRNA primers 27F-YMY and 1492R-HY (Weisburg et al. 1991; Frank et al. 2008; Table S1). Blocking primer sequences were designed from alignments of partial 16S sequences from common bacterial TBPs (e.g., *B. burgdorferi sensu stricto*, *Anaplasma phagocytophilum*) and *R. buchneri,* retrieved from GenBank (Figure 1; Table 1). The forward blocking primer 18F-Rb-C3 and 22F-Rb-C3 overlap the 3′end of 27F-YMY by 9 and 5 nucleotides, respectively, and the 1492R-Rb-C3 overlaps the 5′ end of 1492R-HY by 5 nucleotides. All blocking primer candidates carry a C3 spacer modification (3 hydrocarbons, 1-dimethoxytrityloxy-propanediol-3-succinoyl-long-chain alkylamino) at the 3′ end, preventing oligonucleotide elongation during PCR while exerting minimal or negligible influence on annealing properties (Tan and Liu 2018; Vestheim and Jarman 2008). Blocking primer concentration was empirically optimized to a 7-fold molar excess over each 16S universal primer, thereby selectively suppressing *R. buchneri* amplification while preserving amplification of non-target bacterial taxa (S1 Figure).

**Figure 1:**
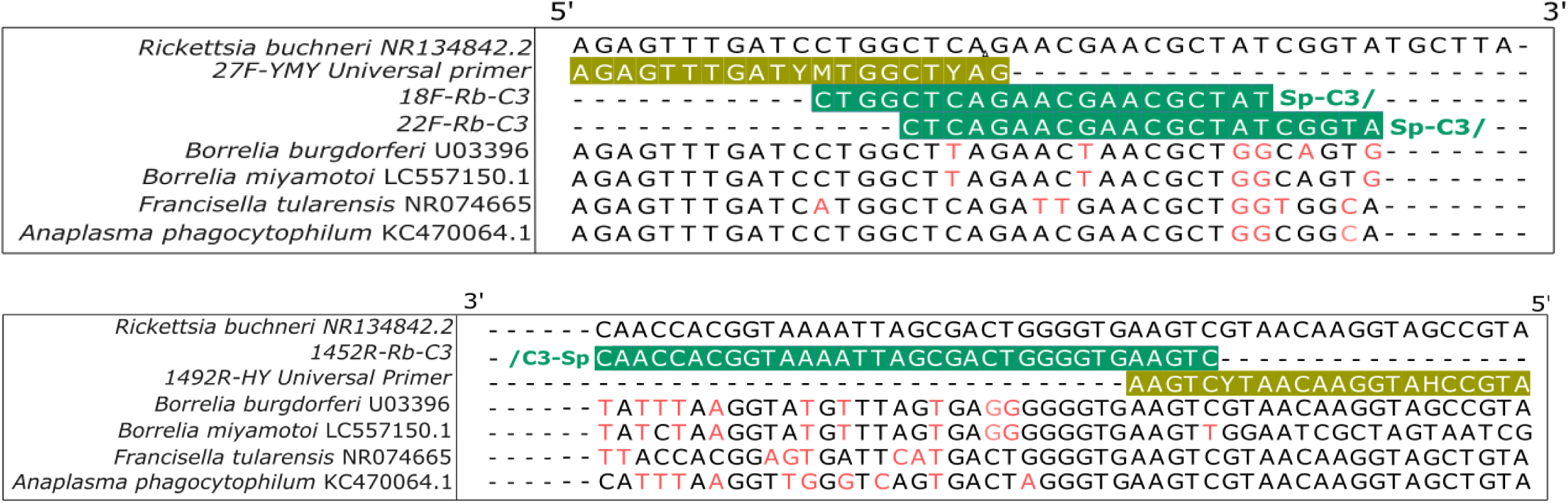
Sequence alignment of blocking primer with 16S rRNA genes from tick-associated bacterial microbiome. The /C3-Sp/ blocking primer (teal highlight) is aligned against 16S rRNA sequences of *Rickettsia*, *Borrelia*, *Francisella,* and *Anaplasma* species, which are commonly present in *Ixodes* species. Mismatched nucleotides between the blocking primers and 16S sequences are highlighted in red. The universal 16S primers are highlighted in olive yellow. All the 16S reference sequences are collected from GenBank.

**Table 1:**
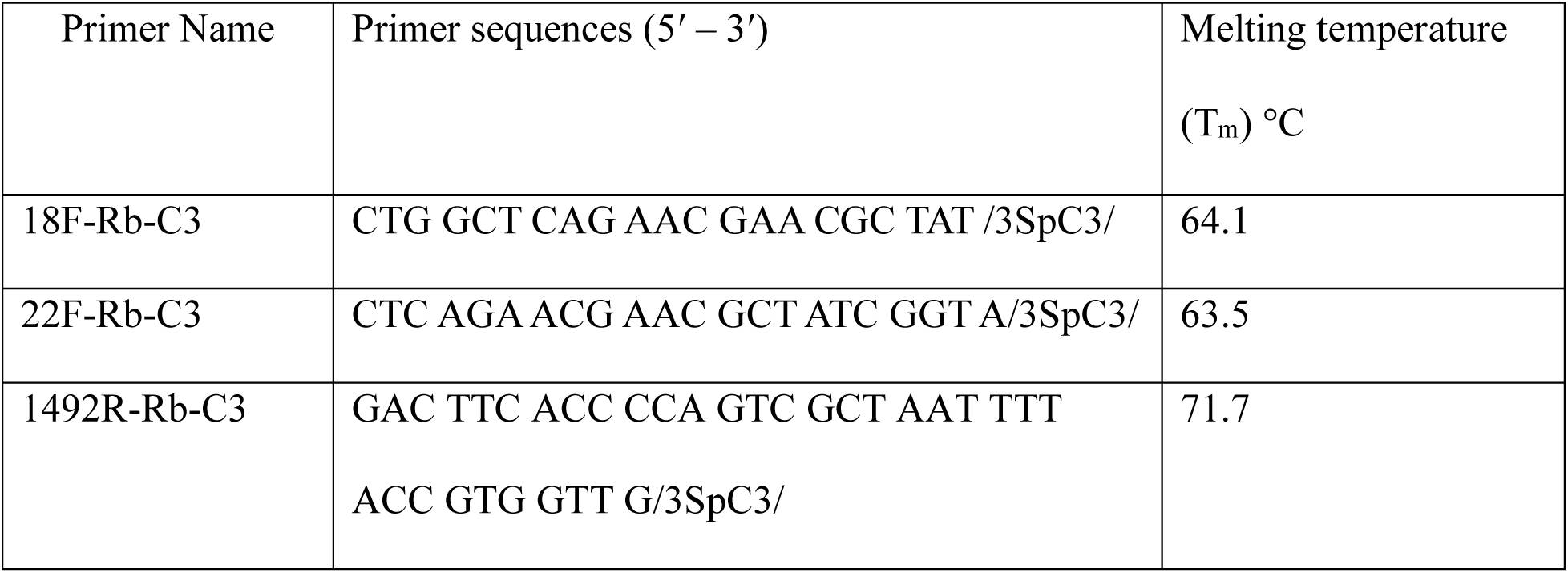
Blocking primers with sequences (5′-3′) and their melting temperature in °C.

### 2.2 Testing and validation of the blocking primers

#### a Sample selection and DNA extraction

Questing adult *Ixodes scapularis* ticks were collected either by flagging or by passive sampling at Queen’s University Biological Station, located 40 km north of Kingston, Ontario, Canada (44.5744°N, 76.3325°W). Ticks were immersed in 2 mL screw-cap tubes containing 70% ethanol, each labeled with a unique barcode generated by baRcodeR (Wu and Colautti 2026), and stored at −20 °C until DNA extraction.

Ticks were surface-sterilized with 5% sodium hypochlorite for 5 minutes and washed three times (3 minutes each) with nuclease-free water to remove the external microbial contamination before DNA extraction. Genomic DNA was extracted using the Cetyltrimethylammonium Bromide (CTAB) protocol, as described in Paulson et al (2023). Briefly, ticks were air-dried for 2 minutes and transferred to sterile 2 mL tubes containing low-binding SPEX ZrO beads (Froggobio, Canada) of sizes 0.2 mm and 0.5 mm in equal volumes to facilitate cell lysis and exoskeleton disruption, respectively. Samples were frozen in liquid nitrogen for 3 minutes before pulverization at 100 Hz for 5 minutes (Next Advance Bullet Blender Storm, USA). 500 μL of preheated CTAB buffer (100 mM Tris-HCl [pH 8.0], 25 mM EDTA, 1.5 M NaCl, 3% CTAB, 1% polyvinylpyrrolidone, and 1% β-mercaptoethanol) was added to pulverized samples, and the tubes were incubated at 62 °C for 16 hours. 500 µL of chloroform was added to each tube, and the tubes were centrifuged at 1500 × *g* for 15 minutes to isolate the genomic DNA. The supernatant was transferred to a 1.7 mL tube, and genomic DNA was precipitated by adding two volumes of ice-cold 100% ethanol. Tubes were mixed by inversion and incubated at −20°C for 30 minutes. Samples were centrifuged at 21,000 × *g* for 15 minutes, washed twice with 1 mL of 75% ethanol, and resuspended in 15 µL of nuclease-free water.

#### b Selection and validation of an effective Blocking primer

A representative tick microbial DNA pool was created by combining 12 individual female adult non-engorged *I. scapularis* DNA extracts and dividing them into four equal-volume aliquots. Aliquots were amplified with 16S rRNA primers 27F-YMY and 1492R-HY; three aliquots each contained one of three blocking primers (18F-Rb-C3, 22F-Rb-C3, or 1492R-Rb-C3), while the fourth was amplified without a blocking primer and served as a control. Amplification was performed in a 50 µL reaction containing 25 µL of 2× PCR Master Mix (FroggaBio, Canada), 2.5 µL each of forward and reverse primers (0.5 µM final concentration), 17.5 µL of blocking primer (3.5 µM final concentration, a 7-fold molar excess over universal primers optimized empirically by gel electrophoresis; S1 Figure), template DNA (15 ng), and nuclease-free water to 50 µL. The reactions were amplified in a thermal cycler (SimpliAmp, Applied Biosystems, USA) under the following conditions: initial denaturation at 95 °C for 5 minutes, followed by 40 cycles of denaturation at 95 °C for 30 seconds, annealing at 55 °C for 30 seconds, and extension at 72 °C for 45 seconds, with a final extension at 72 °C for 10 minutes. PCR products were purified using NucleoMag NGS Clean-up and Size Select Beads (Takara Bio, USA) at a 1.2× volume, then eluted in 10 µL of elution buffer. The amplicon concentration was quantified using a high-sensitivity double-stranded DNA (dsDNA) kit on a DS-11 FX spectrophotometer/fluorometer (DeNovix, USA).

Barcoding and library preparation were performed using the Native Barcoding Expansion Kit (EXP-NBD104) and Ligation Sequencing Kit (SQK-LSK-109; Oxford Nanopore Technologies, United Kingdom). Sequencing was performed on a MinION Mk1C device using a FLO-MIN106D flow cell (R9.4.1 chemistry; Oxford Nanopore Technologies, UK) and managed with MinKNOW v.1.14.1. Raw FAST5 files were converted to FASTQ. Base-calling was performed with Guppy basecaller v.0.5.1 in fast mode, which has lower accuracy but was sufficient to assess blocking primer suppression of *R. buchneri* 16S reads. Downstream analysis was performed using the NaMeco pipeline v.1.2.1 (Yergaliyev et al. 2025), which integrates tools for preprocessing, error correction, and taxonomic classification. Preprocessing was performed using Chopper (De Coster and Rademakers 2023) to retain reads with a minimum length of 1200 bp. Quality-filtered reads were clustered using k-mer counts, followed by dimensionality reduction with UMAP (McInnes et al. 2018) and density-based clustering with HDBSCAN (McInnes et al. 2017), both applied within and between samples using default parameters. Cluster consensus sequences were error-corrected using two rounds of Racon (Vaser et al. 2017) and SPOA (Lee et al. 2002; Lee 2003), and taxonomically annotated using the Genome Taxonomy Database (GTDB. v226; Parks et al. 2020). NaMeco output files were imported into R v.4.4.1 in RStudio (R Core Team 2024; RStudio v.2025.09.0, Posit team 2025) as QIIME2-formatted artifacts and converted into a phyloseq object (McMurdie and Holmes 2013). Community composition was analyzed using microViz v.0.13.0 (Barnett et al. 2021), a wrapper that includes the microbiome (microbiome 2012) and phyloseq. Raw counts were transformed into relative abundance and visualized as a stacked bar plot in ggplot2 v.4.0.2 (Hadley 2016). The OTU table was repeatedly rarefied using vegan v.2.7.3 over 1,000 iterations, and mean values were imported as a phyloseq object. Counts were agglomerated at the species level, and log₂ fold-change values were calculated relative to the control (no blocking primer) and visualized as a heatmap using ComplexHeatmap v.2.22.0 (Z. Gu et al. 2016). Effective blocking primers were assessed by suppression of *R. buchneri* abundance and by the retention of non-target bacterial taxa relative to the control.

The selected blocking primer was validated using 15 individual *I. scapularis* DNA sample extracts (IS1-IS15) and one *Dermacentor variabilis* DNA sample extract (DV-1), included to ensure that the blocking primer did not suppress amplification of closely related non-target taxa. Library preparation and data analysis were performed as described above with the following modifications: barcoded using the Native Barcoding Expansion Kit (EXP-NBD114; Oxford Nanopore Technologies, UK), basecalled using Guppy basecaller v.0.5.1 in default mode and preprocessed to remove reads with a mean PHRED score below 8.

### 2.3 Application of blocking primer in pathogen detection in *I. scapularis*

#### a Sample collection and DNA extraction

67 ticks were collected from farm animals, including bovine, equine, and canine, between 19 May and 23 September 2023 in Eastern Ontario during routine health checks (bovine: n=35; equine: n=22; canine: n=6; questing: n=4). Embedded ticks were placed in 2 mL collection tubes containing 70% ethanol, and each tube was labeled with a unique barcode ID using baRcodeR. Ticks were morphologically identified, transported, and stored at −80 °C until DNA extraction. Ticks were surface-sterilized with 1% sodium hypochlorite for 30 seconds and washed twice with nuclease-free water before extraction. Genomic DNA was extracted as described above.

#### b qPCR analysis

Extracted genomic DNA was screened for *B. burgdorferi sensu stricto* and *B. miyamotoi* using a three-assay quantitative PCR (qPCR) workflow established by the Public Health Agency of Canada (PHAC). First, all samples were screened in triplicate for *Borrelia* spp. targeting the 23S rRNA gene. The samples were deemed presumptive positive with a Ct score < 40 in at least two of the triplicates and were tested for species using a confirmatory assay targeting *ospA* for *B.burgdorferi sensu stricto* and *flaB* for *B. miyamotoi*. The sample was considered positive for the species if the C_t_ score was < 40 in at least two of the triplicates; if none of the triplicates were positive (i.e., C_t_ score = 40), it was categorized as other *Borrelia* spp. Both screening and confirmatory assays were performed, along with positive controls using synthetic DNA and DNA extracted from *B. burgdorferi* cultures, as well as no-template controls. All the assays were performed in a total reaction volume of 25 µL, consisting of 12.5 µL of 2 × Taqman Environmental Master mix (Applied Biosystems, USA), 0.5 µL of each primer with a final concentration of 0.4 µM, 0.15 µL probe with a final concentration of 0.15 µM, 5 µL of template DNA, and 11.35 µL and 10.2 µL nuclease-free water for the 23S rRNA and the *ospA* and *flaB* duplex respectively. The targets were amplified under the following thermocycler conditions (ViiA 7, Applied Biosystems, USA): UNG activation at 50 °C for 2 minutes; 10 minutes of denaturation at 95 °C; 40 cycles of denaturation for 15 seconds at 95 °C; and annealing/extension for 1 minute at 60 °C. The raw data were imported into R v.4.4.1.

#### c Library preparation and Illumina sequencing

Libraries were prepared as described above with modifications summarized here. Samples containing genomic DNA were size-selected using homemade SPRI beads (1.2× volume) to remove DNA fragments shorter than 100 bp. Because the full-length 16S rRNA amplicon exceeds Illumina paired-end read length, a three-step PCR approach was used. PCR1 amplified the full-length 16S rRNA gene using 27F-YMY and 1492R-HY in the presence of the blocking primer to suppress *R. buchneri*. PCR1 product was used as a template for PCR2, which amplified the V4 region of the 16S rRNA gene, suitable for Illumina sequencing. PCR3 added the Illumina indexing adapters to the V4 amplicons.

The reaction of PCR1 was 50 μL, containing 25 μL of Q5 High-Fidelity DNA Master mix (New England BioLabs, USA), 0.4 μM of 27F-YMY and 1492-HY primers, 2.8 μM of 18F-Rb-C3 blocking primer, 3 μL total genomic DNA (∼ 15 to 18 ng), and nuclease-free water to 50 μL. PCR1 was performed in a thermal cycler (T100, Bio-Rad, USA) under the following conditions: initial denaturation at 95°C for 2 minutes, followed by 20 cycles of denaturation at 95°C for 30 seconds, annealing at 62°C for 30 seconds, and extension at 72°C for 1 minute, with a final extension at 72°C for 10 minutes. PCR1 products were purified using SPRI beads at a 1× volume to remove primer dimers and were eluted in 10 µL nuclease-free water.

PCR2 was performed in an 8 µL reaction containing 3 µL of PCR1-purified product and 4 µL of Q5 High-Fidelity DNA Mastermix (New England BioLabs, USA), with 0.5 µL each of the 515F and 806R primers (Apprill et al. 2015; Parada et al. 2016; S1 Table) with spacers as described in Paulson et al. 2023 (S2 Table). Amplification was performed with a thermocycler (T100; Bio-Rad, USA) under the following conditions: initial denaturation at 95 °C for 2 minutes, followed by 20 × cycles of denaturation at 95 °C for 45 seconds, annealing at 55 °C for 45 seconds, and extension at 72 °C for 1 minute, with a final extension at 72 °C for 10 minutes. The PCR2 product was diluted 1:10 with nuclease-free water and used as a template in PCR3.

PCR3 was performed to add the index primers in a 10 µL reaction containing 3 µL of diluted PCR2 product, 1 µL each of i5 and i7 primers, and 5 µL of Q5 High-Fidelity DNA Mastermix (New England BioLabs, USA). Amplification was performed using a thermocycler (T100, Bio-Rad, USA) under the following conditions: initial denaturation at 95 °C for 2 minutes, followed by 10 × cycles of denaturation at 95 °C for 30 seconds, annealing at 65 °C for 30 seconds, and extension for 30 seconds at 72 °C, with a final extension at 72 °C for 10 minutes.

Indexed PCR3 products were normalized using the Just-a-Plate 96 PCR purification and normalization kit (JN-120-10; Charm Biotech, USA) according to the manufacturer’s protocol. Briefly, 10 µL of each indexed PCR3 product was transferred to the normalization plate and mixed with an equal volume of NB7 binding buffer. Plates were sealed, mixed, centrifuged, and incubated at room temperature for at least 1 hour. Wells were washed twice with 50 µL of WB2 washing buffer containing ethanol, air-dried for 10 minutes, and eluted in 20 µL of EB1 elution buffer following a 20 min incubation at room temperature. Normalized libraries were pooled by combining 15 µL from each well into a 1.7 mL tube. An aliquot of 300 µL of normalized pooled DNA was purified using homemade SPRI beads at a 0.8 × bead-to-sample ratio. Beads were incubated with the pooled DNA for 10 minutes at room temperature, washed twice with freshly prepared 80% ethanol, air-dried, and eluted in 20 µL of nuclease-free water. Final pooled libraries were quantified using the NEBNext library quantification kit (New England Biolabs, USA) according to the manufacturer’s protocol. Paired-end sequencing was performed at the Infectious Disease Sequencing Lab in the Kingston Health Sciences Center using the Illumina MiSeq platform with a MiSeq reagent kit v2 (500 cycles; 2 × 250 bp).

#### d Bioinformatic analysis

Raw paired-end reads were demultiplexed into individual FASTQ files using bcl2fastq v.2.2.0 with default settings. Libraries were preprocessed with cutadapt v.5.1 (Martin 2011) to remove primer sequences and low-quality bases (below 20). Reads were further analyzed in R v.4.4.1.

Reads were denoised using the core sample-inference denoising algorithm in DADA2 v.1.30.0 (Callahan et al. 2016). Denoising was performed using default and *prior-*information parameters with pseudo-pooling, incorporating reference 16S rRNA V4 sequences for the Bbsl complex, *B. miyamotoi*, and *A. phagocytophilum,* retrieved from GenBank as *prior* inputs. Forward and reverse reads were trimmed to 210 bp and 200 bp, respectively, clustered into ASVs, and screened for chimeras using the consensus method. Taxonomic assignment of ASVs was performed using the Greengenes2 v.2024.09 (McDonald et al. 2023) naïve Bayes classifier with the DADA2-formatted species-level training set (downloaded 2025; Callahan 2024).

In parallel, preprocessed forward and reverse reads were merged with PEAR v.0.9.11 (Zhang et al. 2014) and converted to FASTA format with seqtk v.1.4 (Li 2012). The merged libraries were taxonomically annotated using BLAST+ v.2.17 (Camacho et al. 2009) against a custom 16S V4 region database comprising sequences of Bbsl, *B. miyamotoi*, and *A. phagocytophilum,* available in GenBank as of 2026, with parameters *e-value < 1e^-20^, percent identity > 97, query coverage > 95, and a maximum of 1 target sequence*. Identical sequences were clustered into BLAST representative sequences, and count and taxonomy tables were constructed using an in-house Bash script. Both tables and the FASTA file were imported into R v.4.4.1, and BLAST representative sequences with fewer than 5 counts or present in only one sample were removed to exclude PCR and sequencing errors.

Samples with an average C_t_ score of 40 were excluded as qPCR-negative before merging the qPCR data with BLAST and DADA2 tables using tidyverse v.2.0.0. The merged file contains reads assigned to various *Borrelia* spp. and their closest species per sample from BLAST, and total ASV count assigned to *B. burgdorferi* and *A. phagocytophilum* per sample in DADA2 under both default and prior information settings. This merged metafile was used to create the Venn diagram, the pie chart, and the UpSet plot using eulerr v.7.1.4 (Larsson and Gustafsson, n.d.), ComplexUpset v.1.3.5 (Krassowski [2020] 2026), and ggplot2 v.4.0.2 (Hadley 2016).

Phylogenetic reconstruction was performed as de with the following modifications. ASVs assigned to *B. burgdorferi* and *A. phagocytophilum* in both the DADA2 default and prior-information datasets, and BLAST representative sequences with the closest hits to *Borrelia* spp., were aligned against a custom 16S rRNA V4 database and reference sequences using DECIPHER v.3.2.0 (Wright 2017). A distance-based tree was constructed as a starting topology, and maximum-likelihood optimization was performed using the best-fit substitution model identified by phangorn v.2.12.1(K. P. Schliep 2011; K. Schliep et al. 2017). Branch support was assessed using 1,000 bootstrap replicates. Final trees were visualized using ggtree v.3.14.0 (Yu et al. 2017), ggtreeExtra v.1.16.0 (Xu et al. 2021), and rphylopic v.1.16.0 (Gearty and Jones 2023).

## 3 Results

### 3.1 18F-Rb-C3 primer suppressed most of the *Rickettsia* reads

Sequencing experiment for screening blocking primer candidates yielded a total of ∼2.9 million reads, with a mean of 0.74 ± 0.34 million reads per sample and a range of 0.32-1.05 million reads. As base calling was performed in fast mode, the PHRED scores were unreliable. Reads were therefore filtered by length rather than by quality score, retaining ∼0.5 million reads with a mean of 0.11 ± 0.04 million reads per sample and a range of 0.05 - 0.15 million reads. Following clustering and error correction in the NaMeco pipeline, 57 consensus clusters were taxonomically annotated using GTDB.

Bacterial community composition at the phylum level showed Pseudomonata dominating most samples (mean relative abundance: 75.79% ± 21.37%; Figure S1). Actinomycetota and Bacteroidota were consistently found across samples, while Bacillota, Bacillota_A, Spirochaetota, and Acidobacteriota were detected in lower abundance. At the species level, *Rickettsia tamurae* was abundant in the sample without blocking primer (relative abundance: 96%), while 25 additional species, including *Cutibacterium acnes* and *Methylobacterium sp. 010692745,* were detected at low abundance (Figure 1A). BLAST alignment against the NCBI rRNA database confirmed that the sequence assigned as *R. tamurae* by Greengenes2 corresponds to the endosymbiont *R. buchneri.* Samples with blocking primers had substantially fewer *Rickettsia*-assigned reads, but the extent of suppression varied among primers. The 18F-Rb-C3 primer demonstrated the highest suppression of *R. buchneri* (∼32-fold reduction), which proportionally increased the relative abundance of *Achromobacter insuavis*, *Cutibacterium flaccumfaciens*, *C. acnes*, *B. burgdorferi*, *Ralstonia pickettii,* and *Methylobacterium sp. 010692745* relative to the control (Figure 1B). Compared to the control, 18F-Rb-C3 also revealed 18 novel taxa and enriched 10 taxa (Table S4). In contrast, 22F-Rb-C3 and 1492R-Rb-C3 revealed only 15 and 11 novel taxa, respectively, and enriched 13 and 10 taxa, respectively, and 22F-Rb-C3 suppressed 10 taxa compared to 13 for 18F-Rb-C3. Overall, initial screening identified 18F-Rb-C3 as the most efficient primer for *Rickettsia* suppression with minimal off-target bias.

Sequencing experiment to validate the blocking primer in individual samples yielded a total of 729,947 reads across 16 samples, with a mean of 45,622 ± 61,401 reads per sample and a range of 5,614 to 272,402 reads. Only 191,749 reads were retained after quality filtering, with a mean of 11,984 ± 9,380 reads per sample and a range of 2,433–28,363 reads. Following clustering and error correction, 84 consensus clusters were generated and taxonomically annotated using GTDB. Bacterial community composition at the species level showed that the mean relative abundance of *R. buchneri* across the *I. scapularis* samples was less than 3%. Following rickettsial suppression, *C. acnes*, *Staphylococcus epidermidis*, *Francisella* sp002095075, *Bartonella contaminerans*, *Stenotrophomonas* unclassified, *Achromobacter xylosidans*, *Comamonas testosteroni*, and *Acinetobacter pittii* were detected in higher abundance (Figure 2C). *B. burgdorferi* was detected in several *I. scapularis* samples, and *Francisella* sp. was detected in *the D. variabilis* sample. Together, both initial screening on pooled samples and individual-sample validation identified 18F-Rb-C3 as the most effective blocking primer for reducing dominant rickettsial-assigned 16S reads while retaining detection of non-rickettsial taxa across samples.

**Figure 2:**
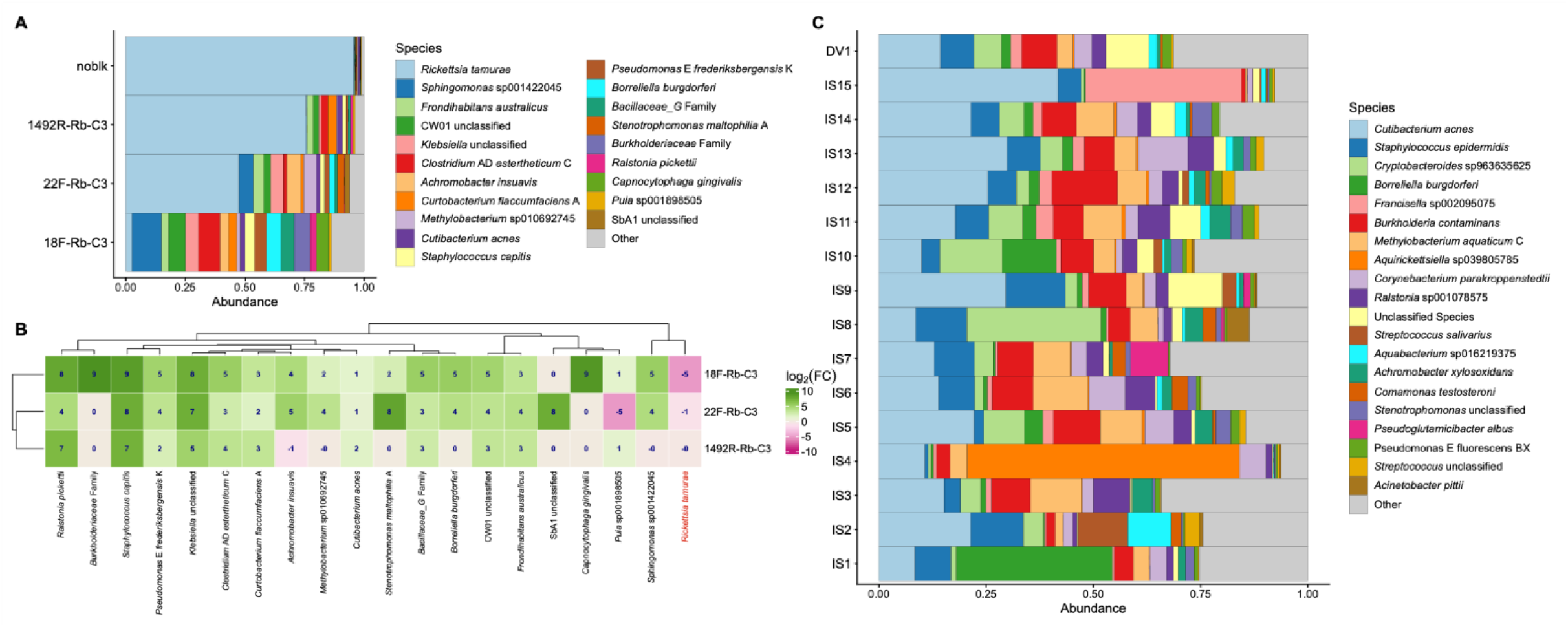
Comparison of *R. buchneri* blocking primers using Nanopore sequencing. **A)** Species-level taxonomic classification of pooled *I. scapularis* DNA extract amplified with full-length 16S rRNA primers comparing species communities across three blocking primers and a sham control. **B)** Heatmap showing Log_2_-fold change in abundance of key bacterial species from panel A. Increases (green) and decreases (purple) in abundance are relative to the control treatment. **C)** Validation of 18F-Rb-C3 blocking primer across individual field-collected questing ticks. Species-level taxonomic classification of 16 questing tick samples, including female adult *I. scapularis* (N=15) and a reference D*. variabilis* (N=1), each amplified with 18F-Rb-C3 blocking primers

### 3.2 qPCR and HTS show complementary detection of *Borrelia* sp. in *I. scapularis* collected from farm animals

A total of 67 ticks were collected, comprising 64 attached to farm animals and 3 questing ticks from Eastern Ontario. Of these, 54 were morphologically identified as adult *I. scapularis*, 11 as adult *Ixodes* spp., and 2 as *D. variabilis*. 61 were female, and six were male; 32 were engorged, and the rest were partially or non-engorged (S5 Table). Only 60 were successfully sequenced; the remaining samples were excluded due to amplification failure. In parallel, 67 genomic DNA extracts were presumptively screened for *Borrelia* spp. by targeting 23S rRNA regions using qPCR.

Sequencing of the 60 viable libraries yielded a total of 14.68 million paired-end reads. 14.60 million reads (99.4%) were retained following quality filtering. Denoising the filtered reads using default parameters in DADA2 generated 2,005 amplicon sequence variants (ASVs), of which only a single variant (ASV46) mapped to *Borrelia* spp. and was detected in 17 samples. Denoising with prior settings in DADA2 yielded 1,895 ASVs, of which only one (ASV45) mapped to *Borrelia* spp. but was detected in 21 samples (S6 Table). In parallel, BLAST-based detection identified 18 samples as positive for *Borrelia* spp. qPCR presumptive screening identified 29 samples as positive for *Borrelia* spp. (S7 Table). Samples that failed sequencing were excluded from qPCR analysis, resulting in 24 samples being positive for *Borrelia* spp. Overall, *Borrelia* spp. detection varied across methods, with qPCR identifying the most positive samples (n=24), followed by DADA2 prior settings (n=21), BLAST (n=18), and DADA2 default settings (n=17).

Comparing the *Borrelia* spp. detection between sequencing (HTS) and qPCR methods showed that 11 samples were deemed positive by both (Figure 3). HTS detected *Borrelia* spp. exclusively in seven samples that were negative in both qPCR replicates, of which six were engorged ticks. Conversely, six samples were missed entirely by HTS but detected by qPCR; four were positive in both qPCR replicates, and the remaining two were positive in one replicate. Three samples were detected by HTS, but only in one of the qPCR replicates. Altogether, these results highlight the complementary nature of the methods, with HTS and qPCR showing partial overlap in *Borrelia* spp. detection across the 60 samples.

**Figure 3:**
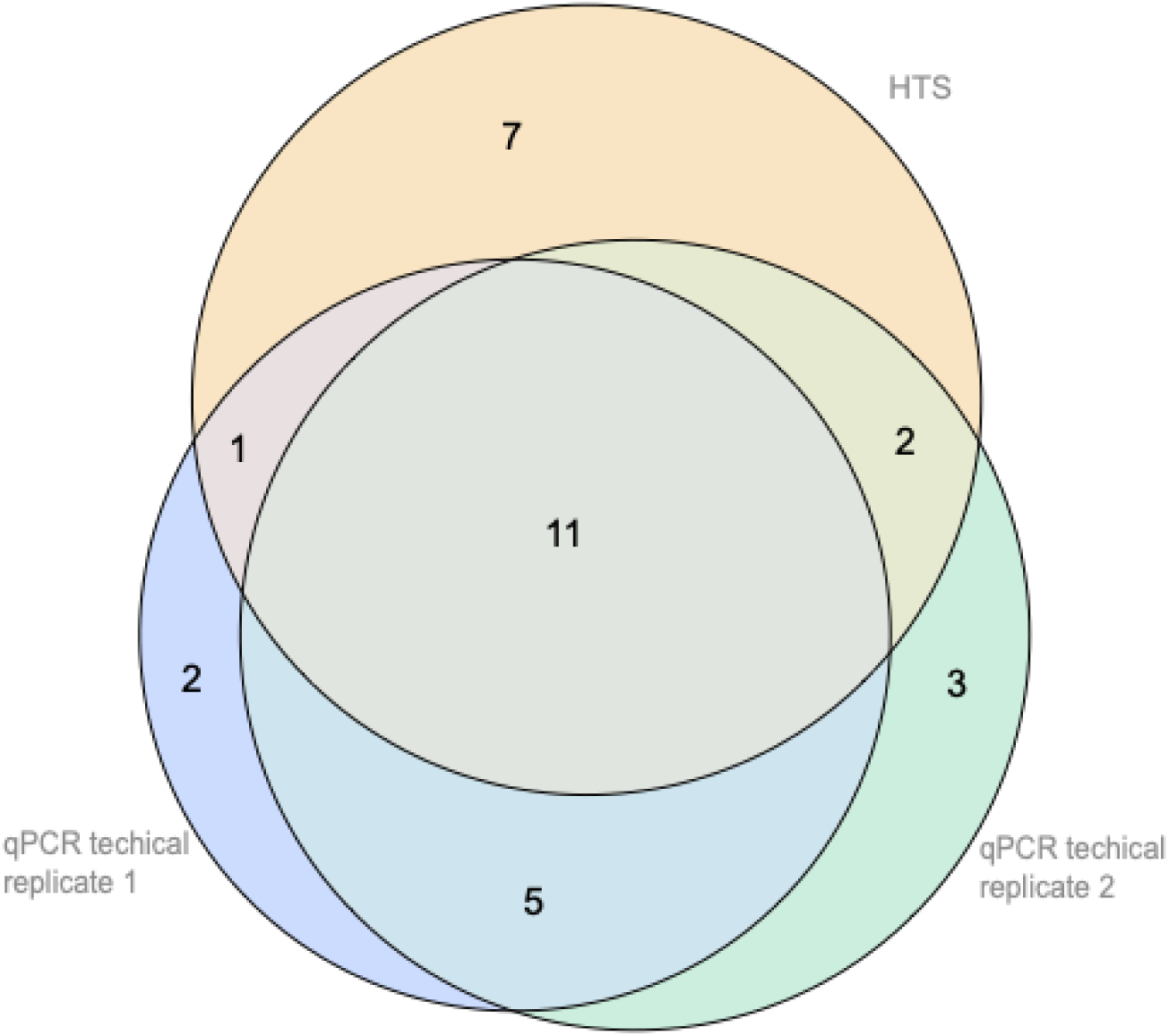
Venn diagram comparing *Borrelia* spp. detection across the same panel of adult *I. scapularis* samples (N=60) collected from farm animals. The number of samples with positive *Borrelia* spp. detection is shown for two qPCR replicates targeting 23S rRNA regions and a single high-throughput sequencing (HTS) analysis of the 16S rRNA V4 region. For example, *Borrelia* was positive in the same 11 samples across all three assays, whereas HTS identified seven samples uniquely positive for *Borrelia* spp., distinct from the five samples positive in both qPCR replicates but not in HTS.

### 3.3 HTS provides agnostic detection of tick-borne bacterial pathogens

Presumptive qPCR-positive samples underwent a secondary qPCR assay targeting *ospA* (*B. burgdorferi* sensu stricto) and *flaB* (*B. miyamotoi*) for species-level confirmation, which identified 11 samples as *B. burgdorferi* sensu stricto, four as *B. miyamotoi*, two as coinfected, and the remaining seven as *Borrelia* spp. (Figure 4a; S7 Table). Of the seven, only three were deemed positive for *Borrelia* spp. as per the PHAC protocol. DADA2 denoising under both default and prior settings collapsed all *Borrelia* variants into a single ASV assigned to *B. burgdorferi* sensu stricto (Figure 4b; S6 Table). In contrast, BLAST-based detection resolved four distinct *Borrelia* variants with more than 97% sequence identity to *B. bissettiae* (NR148750.1), *B. lusitaniae* (NR036806.1), *B. carolinensis* (NR116169.1), and *B. garinii* (NR043413.1), with 5 samples positive for more than one *Borrelia* variant, indicating *Borrelia* co-infection in *I. scapularis*. In addition to *Borrelia* variants, HTS also detected *A. phagocytophilum* in multiple samples under default parameters. Three *Anaplasma*-assigned ASVs were detected in 10 samples: one ASV (ASV13) was detected in all samples, and the re*ma*ining ASVs were detected in only one sample (Figure 5). Under *prior* settings, four *Anaplasma*-assigned ASVs were detected in 12 samples. Overall, these results suggest that while qPCR provides reliable species-level confirmation, HTS captures a broader diversity of TBP agnostically but depends on the parameters used.

**Figure 4:**
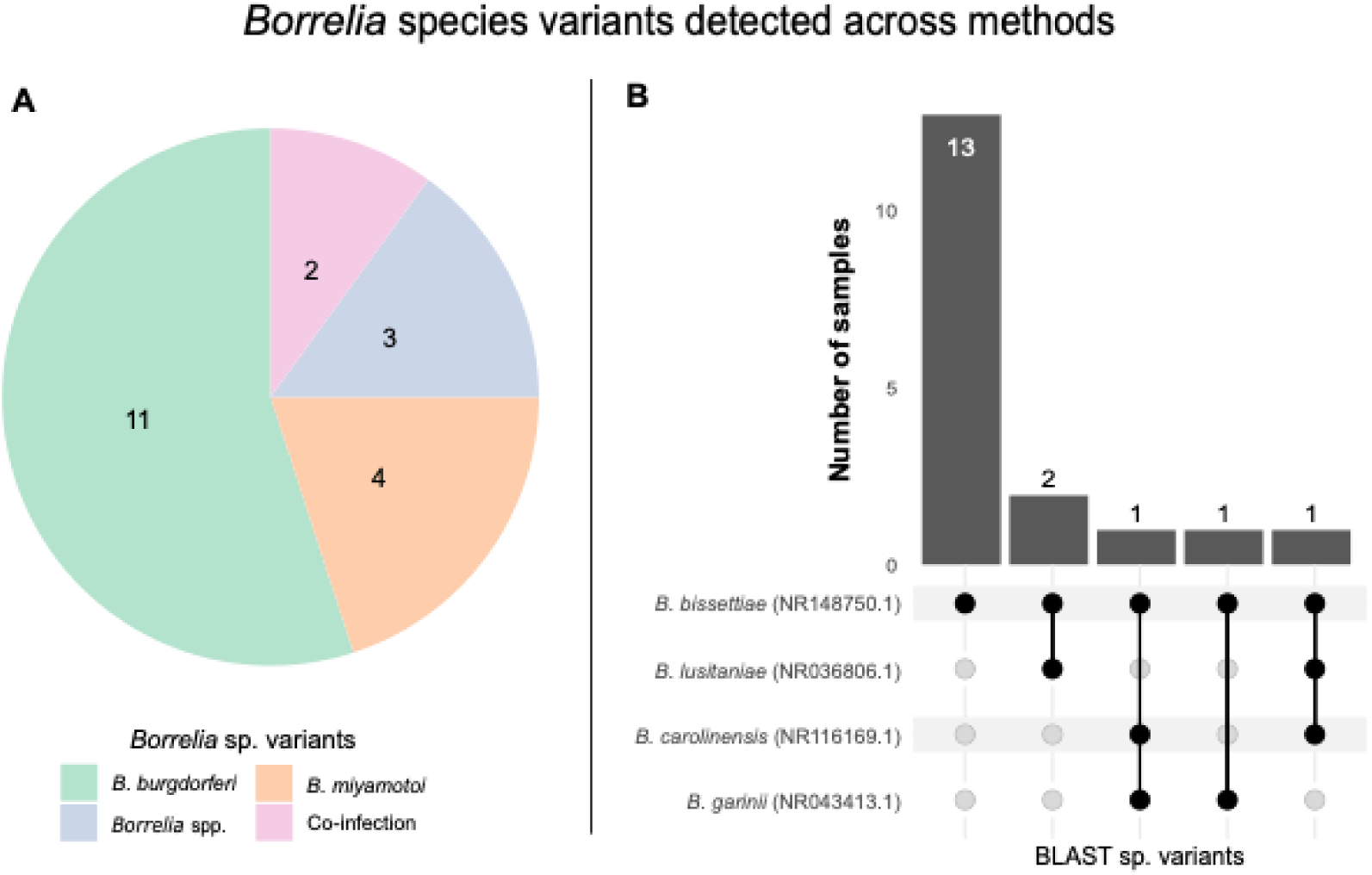
*Borrelia* species and sequence variants detected by qPCR and HTS. (A) Number of samples positive for Borrelia species. in at least one qPCR replicate targeting *ospA* of *B. burgdorferi sensu stricto* (green), *flaB* of *B. miyamotoi* (orange), samples positive for non-specific *Borrelia* 23S rRNA but negative for *ospA* and *flaB* (*Borrelia* spp.; blue), and samples positive for both *ospA* and *flaB* (purple) (B) Species-level resolution using HTS and BLAST. The Upset plot shows *Borrelia* species detected in positive samples (N=18) based on the closest BLAST hit (>97% sequence identity), including NCBI RefSeq accession IDs.

**Figure 5:**
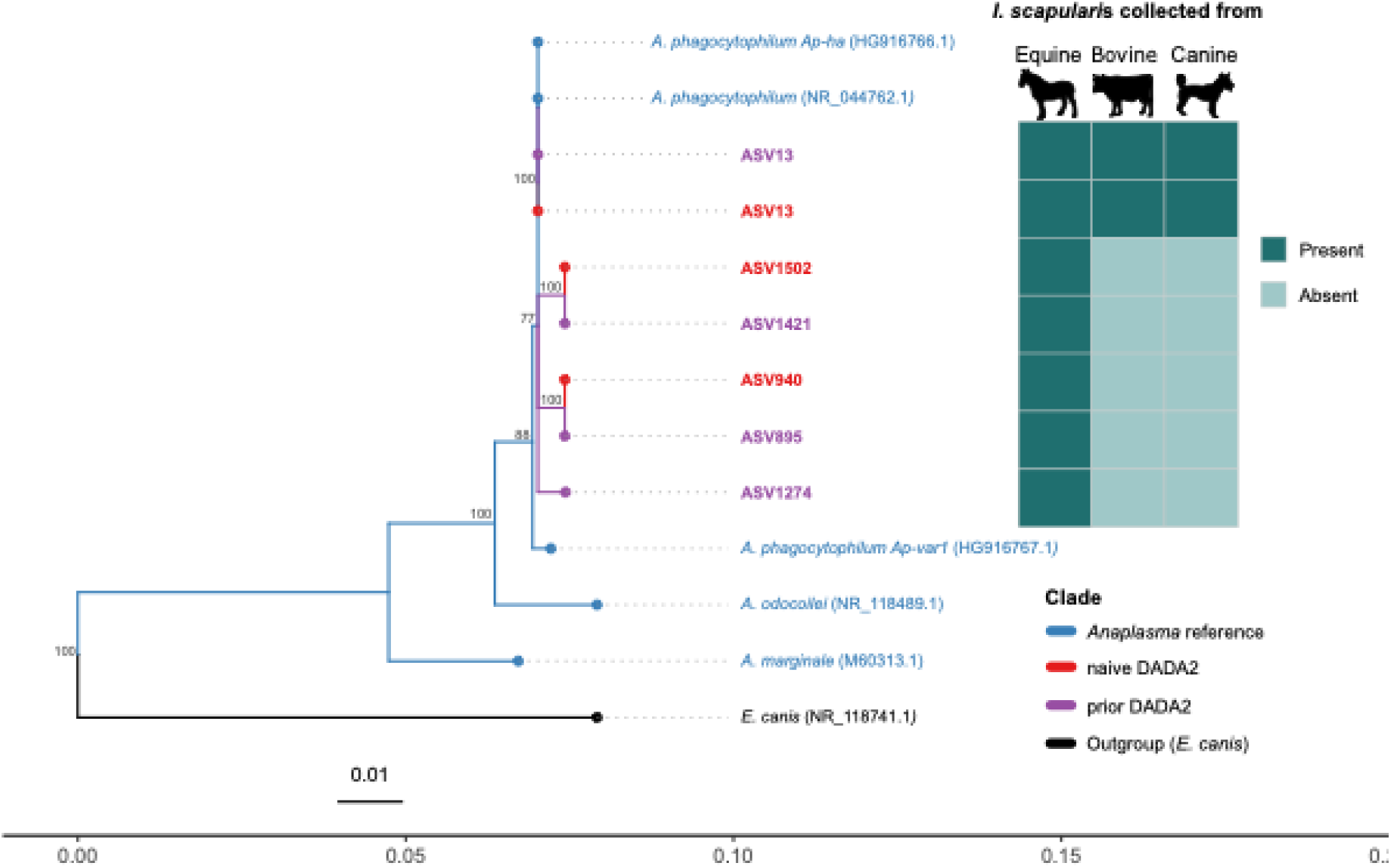
Phylogenetic relationships and host-association heatmap of *Anaplasma* sequence variants detected in *I. scapularis* samples from farm animals. Maximum-likelihood tree of V4-16S rRNA amplicon sequence variants (ASVs) from DADA2 (red and purple) and NCBI RefSeq reference sequences (blue), with *Ehrlichia canis* as the outgroup (black). Branch colors denote ASVs from two DADA2 parameter settings: default (red) and a priori-based (purple). Bootstrap support was assessed using 1000 replicates, with branches > 70% are shown. The heatmap shows the distribution of *Anaplasma* ASVs across *I. scapularis* samples collected from equine, bovine, and canine hosts. Colors in the heatmap indicate the presence (teal) or absence (seafoam green) of each ASV with the host category.

Phylogenetic reconstruction of ASV assigned as *B. burgdorferi sensu stricto* from DADA2 and representative sequences with more than 97% identity across multiple *Borrelia* species variants from BLAST rooted with reference sequences from NCBI RefSeq, placed all within the Bbsl complex (Figure 6). Both representative sequences from DADA2 under default and prior settings (ASV45 and ASV46) grouped closely with *B. burgdorferi* (MH781147.1) and *B. bissettiae* (NR_148750.1), with BLAST Cluster B (N=9) clustering in proximity. In contrast, BLAST Cluster A (N=20) formed a distinct group within the *Bbsl* complex, clustering near *B. carolinensis* (NR_116169.1) with 90% bootstrap support. The relapsing fever group – comprising *B. miyamotoi*, *B. coriaceae*, and several other species formed a well-supported, distinct clade, with no representative sequences clustering nearby. Phylogenetic reconstruction of the ASVs annotated as *A. phagocytophilum* (Figure 5), rooted with *A. marginale* (M60313.1) as the outgroup, placed all sequences within the *A. phagocytophilum* clade, clustering closely with *A. phagocytophilum* (Ap-ha variant) (HG916766.1) and *A. phagocytophilum* (NR_044762.1) with 100% bootstrap support. Overall, 16S HTS, paired with the 18F-Rb-C3 blocking primer to suppress dominant rickettsial reads, enabled broader detection of tick-borne bacterial pathogens than the targeted qPCR assay. However, the V4 region of the 16S rRNA gene lacks sufficient resolution to distinguish closely related bacterial TBP species and strains, and integrating bioinformatic tools with a targeted qPCR assay provides better species resolution.

**Figure 6:**
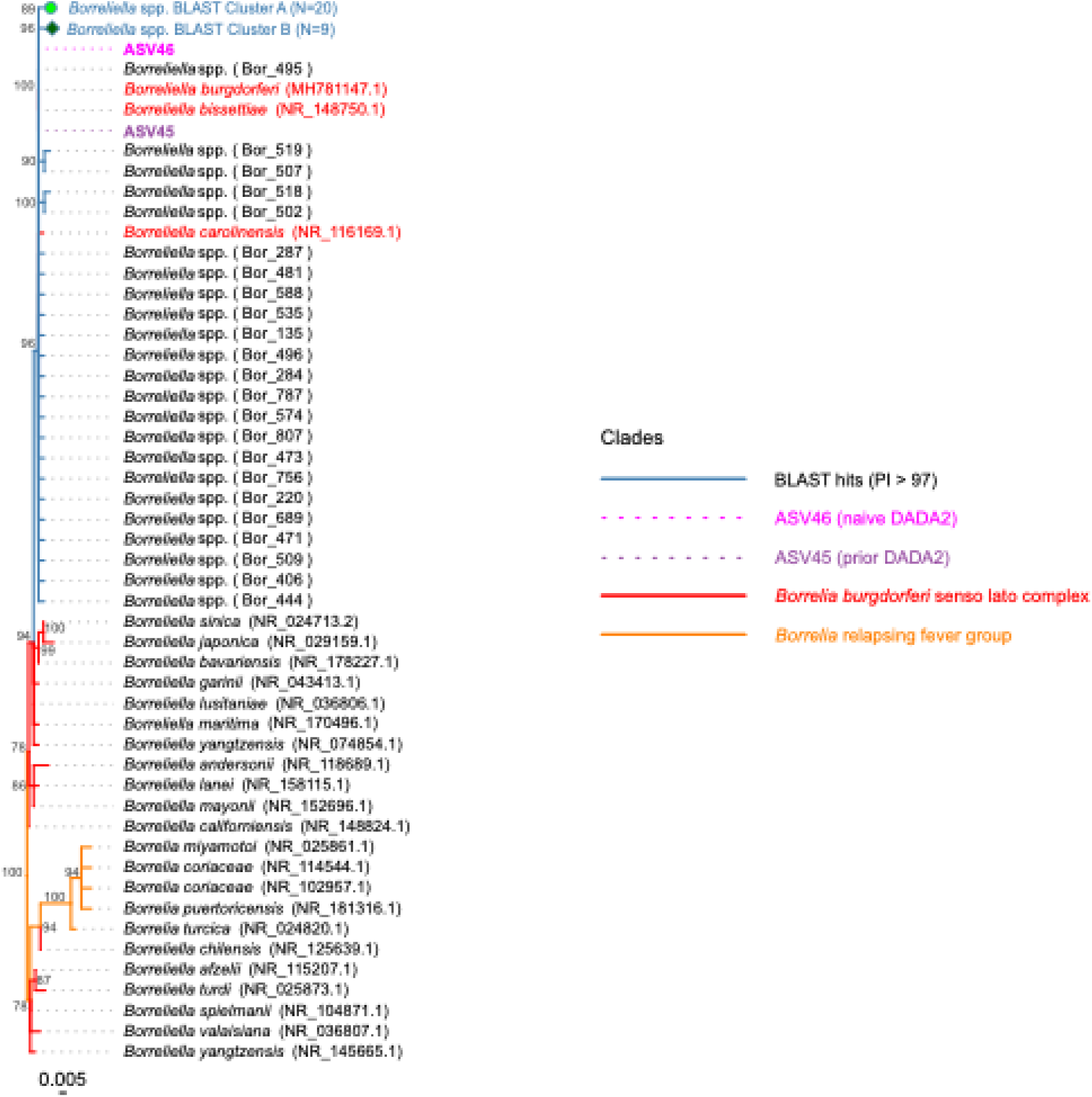
Phylogenetic tree of *Borrelia* spp. detected in *I. scapularis* samples from farm animals. Maximum-likelihood tree of V4-16S rRNA amplicon sequence variants (ASVs) from DADA2 (pink and purple), BLAST-identified sequences (blue), and NCBI RefSeq (red and orange). Branch colors denote two DADA2 parameter settings: default (pink), a priori-based (purple), and BLAST (blue). Bootstrap support was assessed using 1000 replicates, and branches with support >70% were shown.

## 4 Discussion

Here we show that 16S metabarcoding paired with the 18F-Rb-C3 blocking primer improved detection of bacterial TBPs in *I. scapularis*, enabling agnostic detection of bacterial TBPs in a single assay that were undetected by targeted qPCR, which relies on target-specific primers and is susceptible to strain-level sequence variation (Tokarz and Lipkin 2021; Mechai et al. 2025). However, metabarcoding remains more susceptible to detection dropout at lower pathogen loads than targeted qPCR, and detection sensitivity varies across bioinformatic pipelines. Furthermore, the 16S V4 locus lacks sufficient species-level resolution to distinguish closely related *Borrelia* and *Anaplasma* variants. Together, these findings support integrating blocking primer-enhanced HTS with targeted qPCR for comprehensive TBP surveillance.

In an initial screening experiment, we developed three *R. buchneri-*specific annealing-inhibiting blocking primer candidates (18F-Rb-C3, 22F-Rb-C3, and 1492R-Rb-C3; Figure 1) and screened them in pooled I*. scapularis* samples. 18F-Rb-C3 suppressed rickettsial reads by approximately 1,000-fold (Figure 2b) compared to the no-blocking primer control and outperformed 22F-Rb-C3 (relative abundance: 47%; Figure 2a) and 1492R-Rb-C3 (relative abundance: 75%; Figure 2a). The higher blocking efficiency of the 18F-Rb-C3 primer may be partly due to the larger nucleotide overlap of 9 nucleotides with the 27F-YMY universal primer, while 22F-Rb-C3 has only 5 nucleotide overlaps, although blocking efficiency is also influenced by factors such as melting temperature (T_m_), GC content, secondary structure, and primer concentration (Vestheim and Jarman 2008; Tan and Liu 2018). The weaker suppression observed with 1492R-Rb-C3 may reflect self-complementarity or complementarity with the 1492R-HY reverse primer, consistent with previous reports that reverse-blocking primers are prone to self-complementarity-induced reductions in efficiency (Tan and Liu 2018).

Validation of the 18F-Rb-C3 blocking primer in individual field-collected adult female *I. scapularis* ticks confirmed substantial suppression of R. *buchneri*, reducing its mean relative abundance to 3% (SD: ± 8%; Figure 2c). This increased recovery of lower-abundance bacterial taxa, including TBPs such as *Borrelia*, as well as genera often reported as transient or facultative members of the *I. scapularis* microbiome, including *Cutibacterium*, *Staphylococcus*, *Corynebacterium*, *Streptococcus*, *Methylobacterium*, and *Burkholderia*. While such transient taxa are likely acquired horizontally from hosts or the environment and may influence tick immunity and vector competence (Narasimhan and Fikrig 2015; Ross et al. 2018; Narasimhan et al. 2021; Couret et al. 2022). However, such taxa could also result from contamination, as I. scapularis samples tend to have a lower microbial load (Ross et al. 2018). To minimize the risk, we followed stringent clean laboratory practices and, in some experiments, pooled individual tick samples prior to PCR sequencing to obtain a higher microbial load. Overall, the 18F-Rb-C3 blocking primer effectively reallocates sequencing depth from the dominant endosymbiont to the broader bacterial community, improving *I. scapularis* microbiome characterization, including enhanced agnostic TBPs detection.

When blocking primer-enhanced 16S metabarcoding was compared with targeted qPCR, TBPs detection varied across assays and bioinformatic pipelines. Presumptive screening by qPCR targeting the 23S rRNA gene and V4-16S metabarcoding, analyzed with prior-informed DADA2, both identified 11 samples as *Borrelia*-positive (Figure 3). Furthermore, HTS detected *Borrelia*-associated sequences in seven samples that were negative by qPCR. Several mechanisms may plausibly explain the discordance, including reduced primer and probe binding affinity due to sequence variation among *Borrelia* genospecies and strains at the 23S rRNA region (Postic et al. 1994; Margos et al. 2011; Mechai et al. 2025), stochastic detection near the limit of detection, or inhibition of qPCR amplification in engorged ticks (Al-Soud et al. 2000; Al-Soud and Rådström 2001; Dharmarajan and Rhodes 2011). Six of the seven samples exclusively detected by HTS were engorged ticks, consistent with the possibility of blood-associated PCR inhibition (Schrader et al. 2012). However, 23S rRNA target-site divergence and PCR inhibition should both be treated as plausible explanations rather than confirmed mechanisms without direct validation through whole-genome sequencing and inhibition assays, respectively. On the contrary, six samples were positive by *Borrelia* 23S rRNA presumptive qPCR but were not recovered by HTS, suggesting that qPCR had a lower detection threshold for specific pathogens, particularly when pathogen DNA is present at very low abundance. Previous targeted qPCR assays for *B. burgdorferi* reported a limit of detection of 3 to 50 copies per reaction (Courtney et al. 2004a; Graham et al. 2018), whereas 16S metabarcoding depends on both the pathogen template abundance and the proportion of total amplicons represented by the pathogen. Rare and low-abundance taxa are therefore lost during sequencing, library preparation, quality filtering, or denoising (Hatzenbuhler et al. 2017; Cassens et al. 2026). This is a common trade-off between targeted and broad-range surveillance, where targeted assays like qPCR are highly optimized for a predefined panel of TBPs with high sensitivity, while V4-16S metabarcoding provides broader bacterial screening at the cost of reduced reliability for very low-abundance taxa.

Bioinformatic processing also influences TBPs detection. DADA2, under default and *prior* with the pseudo-pooling parameter, detected a single *Borrelia*-assigned ASV. However, in the latter, five additional samples were detected as positive. This difference reflects DADA2’s conservative error model under default settings; rare true variants may fail to pass the statistical threshold for independent inference (Omega A = 1e-40), reducing the recovery of TBP sequences (Callahan et al. 2016). Prior-informed approaches can increase the recovery of rare true variants in target sequences by incorporating known sequence information during inference. This was also evident in the detection of *Anaplasma*, where prior settings recovered one additional ASV that was not detected under default settings (Figure 5). These results show that TBP detection from metabarcoding data is not determined solely by wet-lab performance; it also strongly depends on denoising assumptions, sequence priors, read-depth thresholds, and downstream taxonomic assignment criteria, which are broadly consistent with a recent study on Nanopore Adaptive Sampling (NAS), which had high specificity (0.97) but moderate sensitivity (0.48) for *B. burgdorferi* detection in *I. scapularis*, with the elevated false-negative rate in NAS due to low copy numbers and bioinformatics parameters (Cassens et al. 2026).

Species-level resolution also differed between methods. Targeted qPCR identified ∼ 46% (11/24) as *B. burgdorferi sensu stricto*, ∼17% (4/24) as *B. miyamotoi*, ∼ 8% (2/24) were co-infected with both species, and the remaining ∼29% (7/24) of samples were unresolved, of which only three were confirmed as *Borrelia* spp. positive. *B. miyamotoi* detections had average C_t_ values greater than 39, placing them at the margins of qPCR sensitivity and suggesting an extremely low abundance of *B. miyamotoi* (Table S7). Usually, C_t_ values approaching the maximum threshold of 40 are considered low-confidence detections, as amplification of the target in late cycles is known to be susceptible to technical biases (Kanagawa 2003; Mehle et al. 2014). Therefore, these detections should be interpreted as presumptive even though *B. miyamotoi* has been previously reported in the Ontario region (Grosdidier et al. 2017; Wilson et al. 2023; Crandall et al. 2024), as geographic plausibility alone does not confirm low-template qPCR detection.

In contrast, BLAST-based classification of V4-16S metabarcoding reads resolved four distinct *Borrelia* genospecies. However, these species labels should not be interpreted as confirmed genospecies detection, as V4 regions lack sufficient phylogenetic resolution to reliably distinguish closely related Bbsl lineages (Figure 6), and several of the inferred co-infecting genospecies were represented by fewer than 10 reads (Table S6). These low read counts increase the likelihood that apparent mixed infections may be due to sequencing errors or ambiguous assignments among closely related reference sequences. Genospecies identified in BLAST, including *B. lusitaniae* and *B. garinii*, are commonly associated with European regions (Steinbrink et al. 2022); they should be considered putative. While *B. garinii* has been reported previously in Canadian seabird ticks (*I. uriae*) on coastal islands of Newfoundland and Labrador (Munro et al. 2017), confirmation of this genospecies in Ontario *I. scapularis* would require targeted sequencing of the full-length 16S rRNA gene, multilocus sequence typing (MLST), or whole-genome sequencing.

V4-16S metabarcoding also detected *A. phagocytophilum* in 10 samples under default DADA2 settings and 12 under prior settings; of these, seven samples were coinfected with both *Borrelia* and *Anaplasma*, demonstrating the ability of 16S metabarcoding to detect multiple TBPs in a single assay (Figure S3). However, similar to species resolution of *Borrelia* genospecies, the V4 locus was unable to distinguish between the *A. phagocytophilum* human variant (Ap-ha) and the *A. phagocytophilum* variant 1 (Ap-var) (Figure 5). As *I. scapularis* previously collected in Ontario, Canada may be infected with either *A. phagocytophilum* variant (Krakowetz et al., 2014), with a higher incidence of human granulocytic anaplasmosis (HGA) associated with Ap-ha than with Ap-var1(Prusinski et al. 2023), distinguishing these variants is vital for clinical relevance. Hence, 16S V4 metabarcoding alone cannot provide risk assessment, reinforcing the need for confirmatory genotyping when variant identity affects epidemiological interpretation.

In summary, 16S rRNA metabarcoding paired with the 18F-Rb-C3 blocking primer improved the sensitivity of bacterial TBPs detection in *I. scapularis* by suppressing the dominant rickettsial endosymbiont, thereby enabling agnostic recovery of multiple bacterial TBPs in a single assay. This addresses the key limitation of targeted qPCR, which screens only for certain prevalent TBPs, missing the emerging and novel TBPs, and is vulnerable to sequence variation. However, taxonomic resolution of TBPs is limited by the 16S locus, as V4 regions cannot reliably distinguish closely related genospecies or variants. Robust public health surveillance of TBP using 16S metabarcoding enhanced with a blocking primer should therefore be complemented by targeted assays. This combined strategy preserves the sensitivity and specificity of targeted assays while adding the broader detection capacity needed to identify unexpected, emerging, or previously unscreened bacterial TBPs, thereby aiding risk assessment and policymaking.

## Acknowledgements

We acknowledge the staff and visiting researchers at Queen’s University Biological Station for their assistance with the passive collection of tick samples used in this study. We also acknowledge the staff at Shadow Haven Farms for their assistance in collecting the ticks from farm animals during their routine check. We are also grateful to H. Albers of the Queen’s University Biology Department for their assistance with library preparation, and to the Infectious Disease Sequencing Lab at the Kingston Health Sciences Center (KHSC) for assistance with Illumina sequencing.

S.K.S: Conceptualization, methodology, data curation, visualization, investigation, formal analysis, writing-original draft preparation; S.A.: Conceptualization, methodology, investigation, data curation, writing-reviewing and editing; A.R.P.: Conceptualization, methodology, writing-reviewing and editing; D.B: methodology, writing-reviewing and editing; L.F.C: Resources, funding acquisition, writing-reviewing and editing; J.T: methodology, writing-reviewing and editing; H.W: methodology, writing-reviewing and editing; C.J.S: methodology, writing-reviewing and editing; Shu He: Methodology, writing-reviewing and editing; P.M.S: Methodology, writing-reviewing and editing, funding acquisition; Z.S: Conceptualization, methodology, writing-reviewing and editing; R.I.C: Conceptualization, methodology, investigation, project administration, supervision, funding acquisition, resources, writing-reviewing and editing.

This work was funded by the Canadian Institutes of Health Research - Health Canada’s Healthy Environments and Consumer Safety Branch (CIHR FRN 192227), provided to R.I.C. L.F.C. received grant number 382888 from the Queen’s Faculty Association Fund for Scholarly Research and Creative Work and Professional Development (Adjuncts). P.M.S is supported by and is the recipient of the Jay and Kendal Patry Chair in Clinical Genomics. The funders had no role in study design, data collection, interpretation, or the decision to submit the work for publication.

## Supplementary files

**Figure S1:**
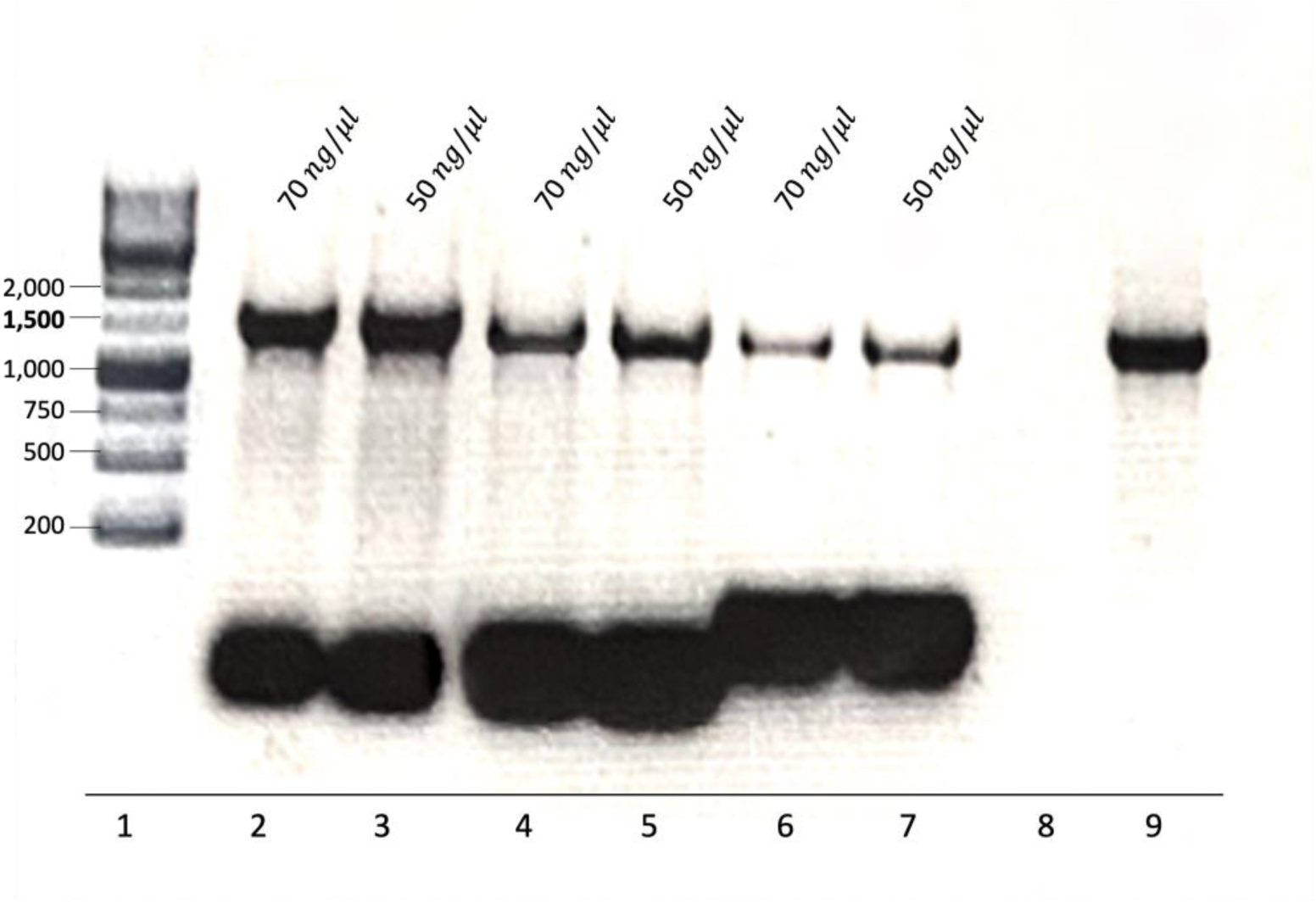
Agarose gel electrophoresis demonstrating the efficacy of *Rickettsia buchneri* - specific blocking. Full-length 16S rRNA PCR was performed using 27F-YMY and 1492R-HY primers in the presence of blocking primers at two concentrations (50 ng/µL and 70 ng/µL). Lane 1:1 kb molecular weight marker; lanes 2–3:1452R-Rb-C3 primer; lanes 4-5: 22F-Rb-C3; lanes 6 and 7:18F-Rb-C3; lane 8: negative control (no template DNA); lane 9, positive control (template DNA without blocking primer).

**Figure S2:**
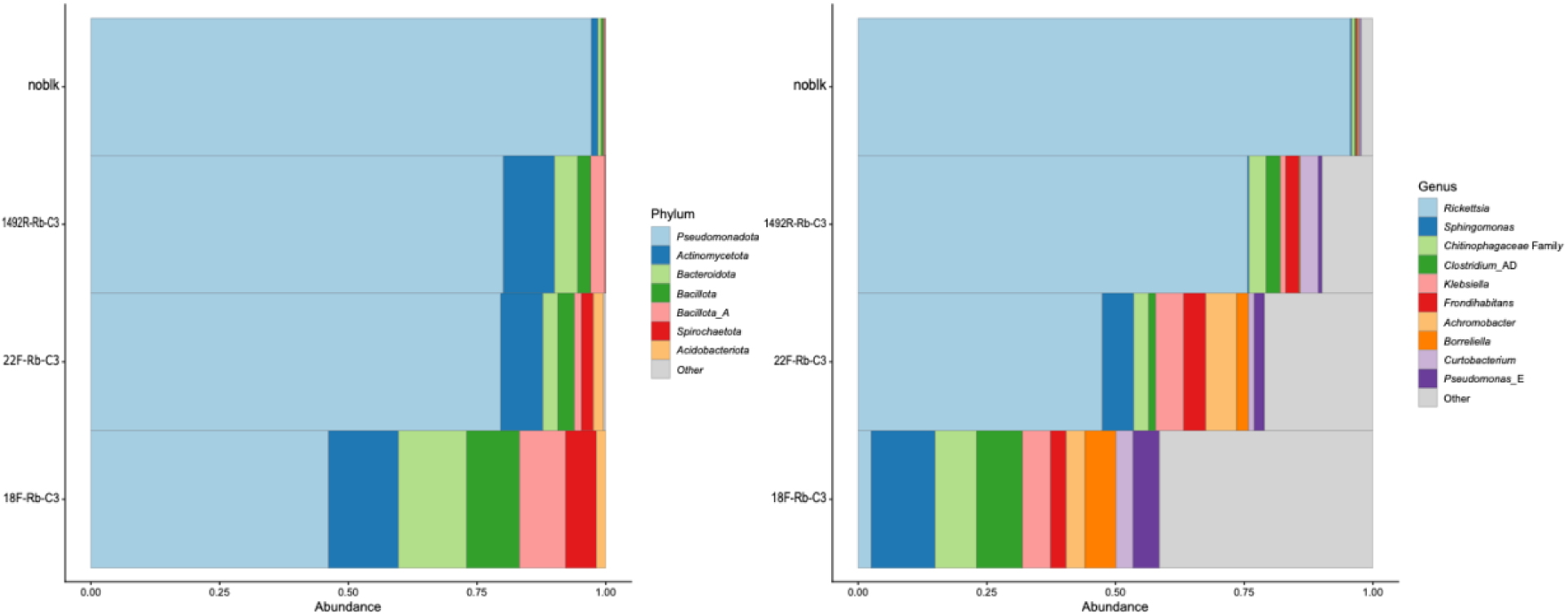
Phylum (left) and Genus (right) level taxonomic classification of a pooled *I. scapularis* sample tested with different blocking primers, along with a no-blocking-primer control. The stacked bar plot shows the relative abundance of the top 7 phyla and the top 10 genera across all samples.

**Figure S3:**
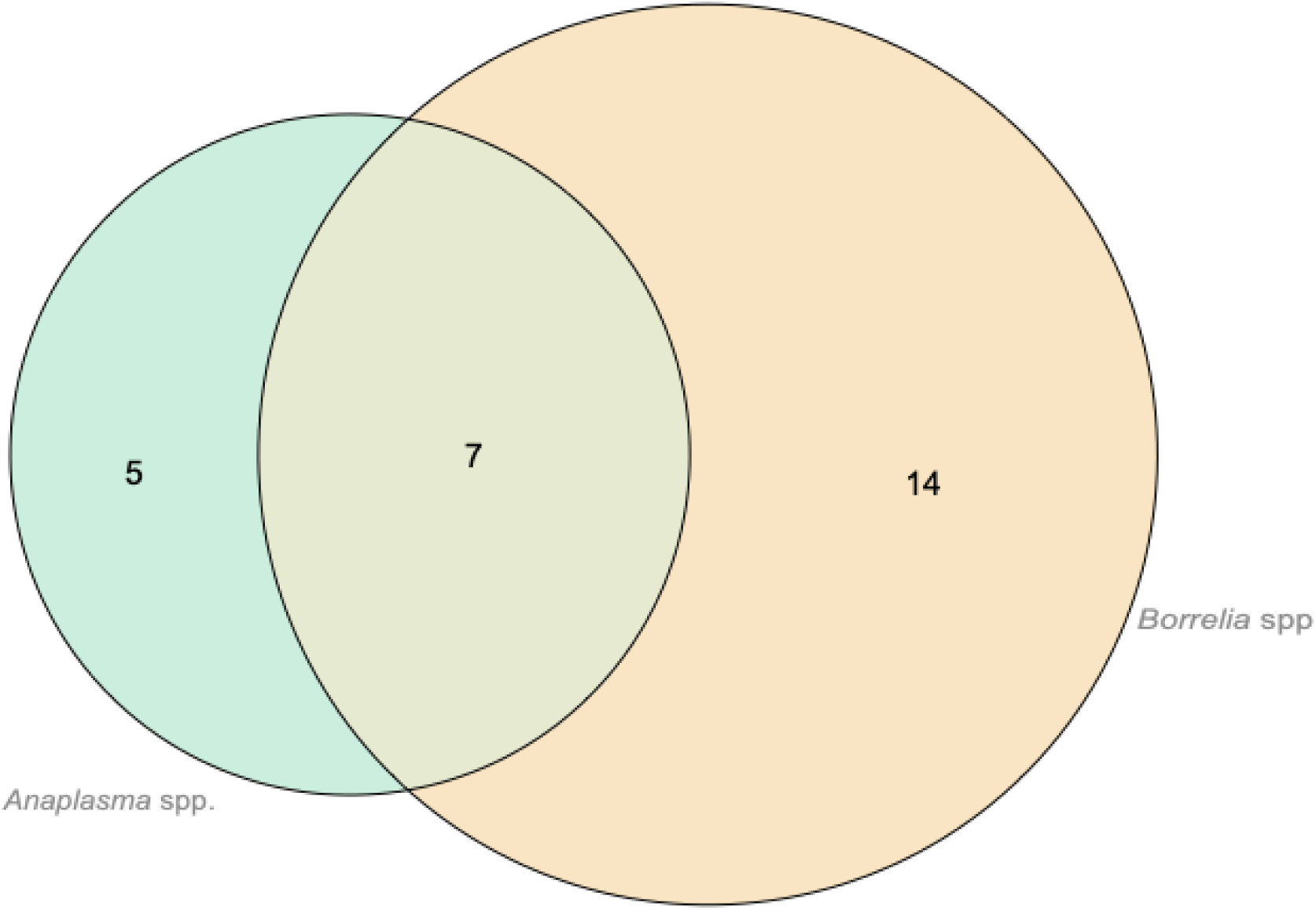
Venn diagram comparing bacterial TBP coinfection in the same panel of adult *I. scapularis* samples (N=60) collected from farm animals. Number of samples positive for Borrelia species and for Anaplasma species in a single high-throughput sequencing (HTS) analysis of the V4 region of 16S rRNA. For example, seven samples were coinfected with both *Anaplasma* spp. and *Borrelia* spp., 14 were positive only for *Borrelia* spp., and five were positive only for *Anaplasma* spp.

---

**Table S1:**
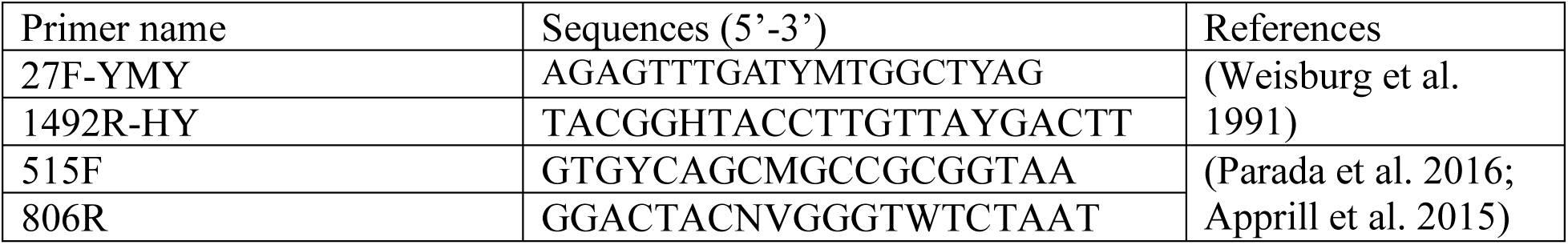
16S full-length and V4 region primers used for targeted amplification, along with their references.

**Table S2:** 16S V4 region primers (515F-806R) with spacers, sequencing primers, composite primer, and indexing primers used in the Illumina sequencing experiment.

**Table S3:**
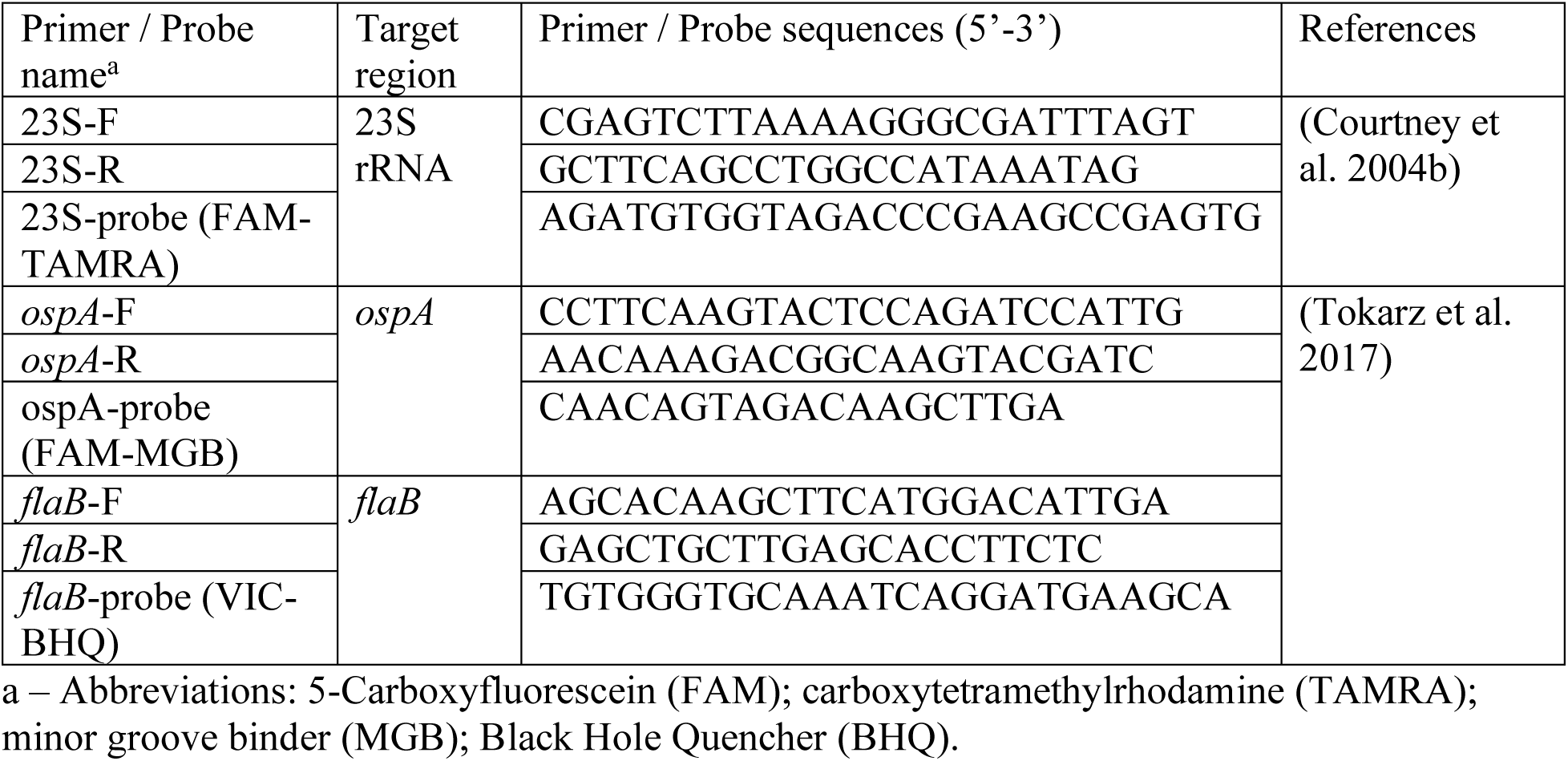
quantitative PCR (qPCR) primers and probes targeting 23S rRNA (*Borrelia* spp.), *ospA* (*B. burgdorferi sensu stricto*), and *flaB* (*B. miyamotoi*) with their references.

**Table S4:** Number of consensus clusters suppressed, reduced, unchanged, enriched, or novel across samples amplified with different blocking primers relative to the no-blocking-primer control sample.

| Blocking primer name | Suppressed | Reduced | Unchanged | Enriched | Novel |
| --- | --- | --- | --- | --- | --- |
| 18F-Rb-C3 | 13 | 1 | 3 | 10 | 18 |
| 1492R-Rb-C3 | 11 | 1 | 5 | 10 | 11 |
| 22F-Rb-C3 | 10 | 1 | 3 | 13 | 15 |

**Table S5:** Metadata of ticks attached to farm animals Should upload the entire sheet

**Table S6:** *Borrelia* species detection from ticks collected from the farm animals using DADA2 and BLAST

**Table S7:** quantitative PCR (qPCR) scores of both screening (23S rRNA for *Borrelia* spp.) and confirmatory (*ospA* for *B. burgdorferi sensu stricto* and *flaB* for *B. miyamotoi*) assays.

## Data availability

The raw data generated in the sequencing experiments have been uploaded as FASTQ data to the GenBank Sequence Read Archive (SRA). Codes and scripts used to generate the results will be uploaded to GitHub for reproducibility.

